# A critical role for E2-p53 interaction during the HPV16 life cycle

**DOI:** 10.1101/2021.11.01.466861

**Authors:** Christian T. Fontan, Claire D. James, Molly L. Bristol, Apurva T. Prabhakar, Raymonde Otoa, Xu Wang, Elmira Karimi, Pavithra Rajagopalan, Devraj Basu, Iain M. Morgan

## Abstract

Human papillomaviruses (HPV) are causative agents in ano-genital and oral cancers; HPV16 is the most prevalent type detected in human cancers. The HPV16 E6 protein targets p53 for proteasomal degradation to facilitate proliferation of the HPV16 infected cell. However, in HPV16 immortalized cells E6 is predominantly spliced (E6*) and unable to degrade p53. Here we demonstrate that human foreskin keratinocytes immortalized by HPV16 (HFK+HPV16), and HPV16 positive oropharyngeal cancers, retain significant expression of p53. In addition, p53 levels can be increased in HPV16+ head and neck cancer cell lines following treatment with cisplatin. Introduction of full-length E6 into HFK+HPV16 resulted in attenuation of cellular growth (in hTERT immortalized HFK, E6 expression promoted enhanced proliferation). An understudied interaction is that between E2 and p53 and we investigated whether this was important for the viral life cycle. We generated mutant genomes with E2 unable to interact with p53 resulting in profound phenotypes in primary HFK. The mutant induced hyper-proliferation, but an ultimate arrest of cell growth; β-galactosidase staining demonstrated increased senescence, and COMET assays showed increased DNA damage compared with HFK+HPV16 wild type cells. There was failure of the viral life cycle in organotypic rafts with the mutant HFK resulting in premature differentiation and reduced proliferation. The results indicate that the E2-p53 interaction is critical during the HPV16 life cycle, and that disruption of this interaction has anti-viral potential. We discuss potential mechanisms to explain these phenotypes.

**Importance:** Human papillomaviruses are causative agents in around 5% of all cancers. There are currently no antivirals available to combat these infections and cancers, therefore it remains a priority to enhance our understanding of the HPV life cycle. Here we demonstrate that an interaction between the viral replication/transcription/segregation factor E2 and the tumor suppressor p53 is critical for the HPV16 life cycle. HPV16 immortalized cells retain significant expression of p53, and the critical role for the E2-p53 interaction demonstrates why this is the case. If the E2-p53 interaction is disrupted then HPV16 immortalized cells fail to proliferate, have enhanced DNA damage and senescence, and there is premature differentiation during the viral life cycle. Results suggest that targeting the E2-p53 interaction would have therapeutic benefits, potentially attenuating the spread of HPV16.

## Introduction

HPV16 infection causes half of cervical cancers and up to 90% of HPV-positive oropharyngeal cancers (1). Despite advances in vaccination, the prevalence in HPV-associated oropharyngeal cancer continues to rise, contributing to an ongoing public health crisis without any effective anti-viral therapies (2–5).

HPV infects basal epithelial cells and delivers its circular 8kbp DNA genome into the nucleus of the host. Consequently, cellular host factors initiate transcription from the viral long control region (LCR) (6). The viral mRNA is expressed from a single transcript which is then processed, spliced and translated into individual viral proteins. In high-risk HPV infection, the viral oncoproteins E6 and E7 contribute to cellular transformation and cancer progression by targeting several cellular proteins, including tumor suppressors p53 and pRb respectively (7–11).

HPV uses two proteins to initiate replication of the viral episome. The E2 protein homodimerizes via a carboxyl terminus domain and binds to four 12-bp palindromic sequences within the viral LCR and origin (12). Via its amino terminus, E2 recruits the viral helicase E1 to the origin which forms a di-hexamer that replicates the viral genome with the assistance of host polymerases (13, 14). Upon initial infection, the HPV genome replicates to 20-50 copies and maintains this copy number as the infected cell migrates through the epithelium. As the infected cell differentiates and reaches the upper layers, the viral genome amplifies and expresses the L1 and L2 capsid proteins that promote virus assembly and shedding (15–17).

E2 has additional roles in the viral life cycle. E2 can promote or repress viral transcription depending on protein concentration (18). E2 binds to host mitotic chromatin to facilitate viral segregation, resulting in sorting of viral episomes into daughter nuclei following mitosis (19). E2 also regulates expression of host genes important in the viral life cycle and cancer progression (20–24).

The tumor suppressor p53 primarily functions as a transcription regulatory factor during cellular stress and DNA-damage, leading to cell cycle arrest, senescence, and apoptosis (25–30). In HPV infection, the 150 amino acid E6 oncoprotein interferes with the transcriptional activity of tumor suppressor p53, as well as induces its degradation (7, 9, 31). Direct degradation is initiated by the formation of a complex with p53 and the host partner protein E6AP, which is an ubiquitin ligase. E6 directs the ligase activity of E6AP to p53, promoting its degradation via the proteasome (7, 31–33). This is in direct contrast with many HPV-negative cancers where p53 often becomes mutationally inactivated (34, 35). The degradation of p53 by HPV is also regulated by alternative splicing of high-risk E6 proteins, resulting in a short modulatory isoforms of the oncoprotein such as E6* and E6*I which do not bind to p53 and can inhibit E6-E6AP-p53 complex formation preventing p53 degradation in a cell-cycle specific manner (36–38). We, and others, have reported that expression of alternatively spliced forms of E6 are more dominant compared to full-length E6 in HPV-positive head and neck cancer, and ectopic expression of these isoforms have anti-tumorigenic effects (38–41). The disruption of the E6-E6AP-p53 degradation complex by E6* and E6*I, allowing for p53 expression, suggests that p53 expression may be important in HPV immortalized cells.

Another reported viral interactor with p53 is E2. E2 proteins from high-risk HPVs can bind to p53 and this interaction can trigger apoptosis in cervical cancer cells (42, 43). Additionally, E2 replication function is regulated by this interaction with p53 as overexpression of p53 reduces viral replication (44). This led us to hypothesize that in HPV16 immortalized cells, residual p53 is necessary to maintain a healthy viral life cycle, perhaps via an interaction with E2.

Here, we demonstrate that p53 expression is clearly detectable in a variety of cell lines and tumors immortalized by HPV16 (45, 46). Depletion of residual p53 by overexpression of full length E6 protein led to a significant reduction in cellular proliferation in keratinocytes immortalized by wild-type HPV16. Expression of a mutant E6 protein lacking p53 binding had no effect on cell growth.

To determine whether an interaction between E2 and p53 was important for cellular proliferation of HPV16 positive cells, we generated an HPV16 genome with point mutations in the E2 gene eliminating p53 binding (42, 43). Keratinocytes were then immortalized by this mutant. Remarkably, compared to wild type HPV16, E2^−p53^ mutant cells, following an initial burst in proliferation, stopped growing and had increased levels of senescence, and accumulated DNA breaks as evidence by single-cell gel electrophoresis assay (COMET). When subjected to differentiation via organotypic raft culturing, these mutant cells had reduced proliferation leading to marked reduction in raft thickness. There was also a reduction in viral replication markers in the mutant cells. These results suggest that although p53 is downregulated by E6 in high-risk HPV infection, p53 is still necessary to permit HPV induced proliferation and that the interaction with E2 plays an important role in the requirement for p53 expression.

## Results

### Tumor Suppressor p53 is expressed in HPV16 immortalized cells and is critical for their optimal growth

Previous studies demonstrated that alternative splice variants (E6*) are the dominant E6 transcripts in HPV associated head and neck cancer, preventing E6-E6AP-p53 complex formation and inhibiting p53 degradation (36–41). We confirmed the presence of p53 in a series of HPV16 positive cell lines (Fig. 1). Expression of the entire HPV16 full genome in N/Tert-1 cells results in partial reduction in p53 compared to near complete abrogation by expression of HPV16 E6 and E7 (Fig. 1A compare lanes 2 and 4 to lane 1). Moreover, human tonsil cells immortalized by HPV16 retain p53 expression near N/Tert-1 control cells (compare lane 1 to 3). To further investigate these findings, we studied two independent donors of human foreskin keratinocytes (HFK) immortalized with HPV16. In both donor lines, p53 levels were much less reduced compared to HFK immortalized by HPV16 E6 and E7 overexpression (Lanes 5-8). To determine whether this expression is affected by tumor microenvironment, we surveyed p53 expression in 8 patient derived xenografts (PDX) from oropharyngeal and oral cavity carcinomas (four HPV16 positive and four negative) (45, 46). All HPV16 positive PDX samples and 3 out of 4 HPV negative retained detectable p53 expression illustrating no clear association between HPV status and p53 expression (Fig. 1B). Platinum based DNA-damaging agents such as cisplatin are critical in the treatment of late stage systemic head and neck cancers (47–50). Because DNA-damage is known to stabilize and activate p53, and p53 is most often wild-type in HPV-positive cancers, we predicted that in HPV+ head and neck cancer cell lines the expression of active wild-type p53 can be promoted by cisplatin treatment. We confirmed dose-dependent cisplatin induced p53 expression in SCC-47 and SCC-104 cells (Fig. 1C). These results indicate that p53 expression is conserved in many cell lines immortalized by the HPV16 genome and can be induced by treatment with DNA-damage. This suggests that although E6 degrades p53 to help promote cell immortalization and carcinogenesis, p53 often retains expression under a variety of conditions, indicating that it still may play an important role in the HPV16 life-cycle.

**Figure 1.**
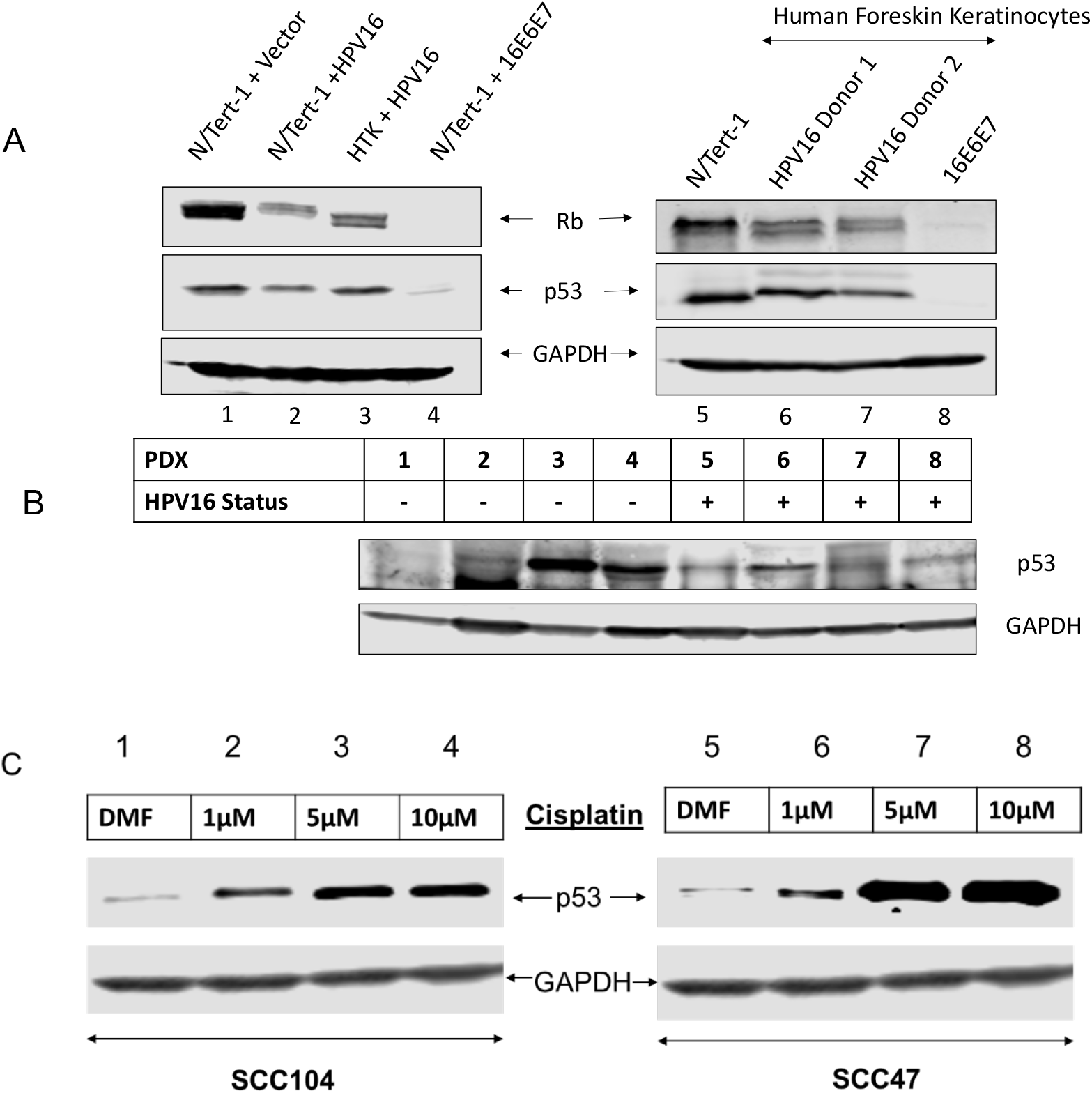
Tumor suppressor p53 expression is conserved in HPV16 immortalized cell lines and patient derived xenografts. (A) Western blot analysis of p53 and pRb in N/tert-1 cells stably expressing the HPV16 genome (lane 2), HPV16 E6+E7 (lane 4) or empty vector (lane 1) compared to Human tonsillar keratinocytes immortalized with HPV16 (lane 3). Two independent human foreskin keratinocyte (HFK) donors were immortalized with wild-type HPV16 (lanes 6 and 7) or overexpression of HPV16 E6 and E7 oncogenes (lane 8, cells from donor 1). (B) Western blot analysis of p53 expression in 4 HPV-negative and 4 HPV-positive patient derived xenografts. (C). p53 expression following 24-hour cisplatin treatment of primary HPV-positive head and neck cancer cell lines (SCC47 and SCC104) DMF solvent only was used as drug-free control. All western blots utilized GAPDH as an internal loading control.

To determine whether reduction of p53 would compromise the growth of HPV16 immortalized cells we introduced full length E6 (using a retroviral delivery of the E6 gene which does not allow alternative splicing) into N/Tert-1 (foreskin keratinocytes immortalized by telomerase) and HFK+HPV16 cells. Fig. 2A demonstrates that the additional expression of E6 in N/Tert-1 cells results in significantly increased cellular proliferation as has been described (51). However, introduction of E6 into HFK+HPV16 resulted in an attenuation of cell growth (Fig. 2B). Because E6 possesses a variety of mechanisms for regulating cellular proliferation independent from p53 degradation, we attempted to isolate these other mechanisms by expressing an E6 mutant unable to promote degradation of p53 but retains all other known functions (52). This mutant did not have a deleterious effect on cell growth indicating that it is the E6 targeting of p53 that results in the attenuation of cellular proliferation. Additionally, we found that these proliferation rates were inversely correlated with senescence levels (Fig. 2C and 2D). In the HFK+HPV16+E6 cells, we noticed that over time the cells began proliferating once again. To determine whether the recovered cells had a restoration of p53 protein levels we carried out western blots of HFK+HPV16+E6 cells at different stages following E6 introduction (Fig. 2E). Lane 4 demonstrates that there is an initial reduction in p53 protein levels in these cells immediately following selection compared with control cells (compare lane 4 with lane 3). However, following 13 days of culturing (when we noticed proliferation begin to restore to that of the control cells) there is a restoration of p53 protein expression (compare lane 7 with lane 4). These results suggest that reduction of p53 protein may lead to growth attenuation and enhanced senescence of HFK+HPV16 cells. They also suggest that to begin to proliferate again, restoration of p53 likely helps promote growth in the HFK+HPV16+E6 cells. We monitored the exogenous E6 RNA levels (Fig 2F). There is a clear reduction in the E6 RNA expressed from the exogenous vector between days 0 and 13 correlating with the restoration of p53 protein expression and cellular proliferation. When we analyzed for E6 protein expression via western blot, while we did not notice an appreciable change in E6 protein levels, we found that there was a significant increase of E6Δp53 expression compared to wild-type E6 at both time points (Fig. 2G). This supports our claim that expression of E6 is more deleterious to growth than E6Δp53 in HFK+HPV16 and this is most likely due to p53 degradation.

**Figure 2.**
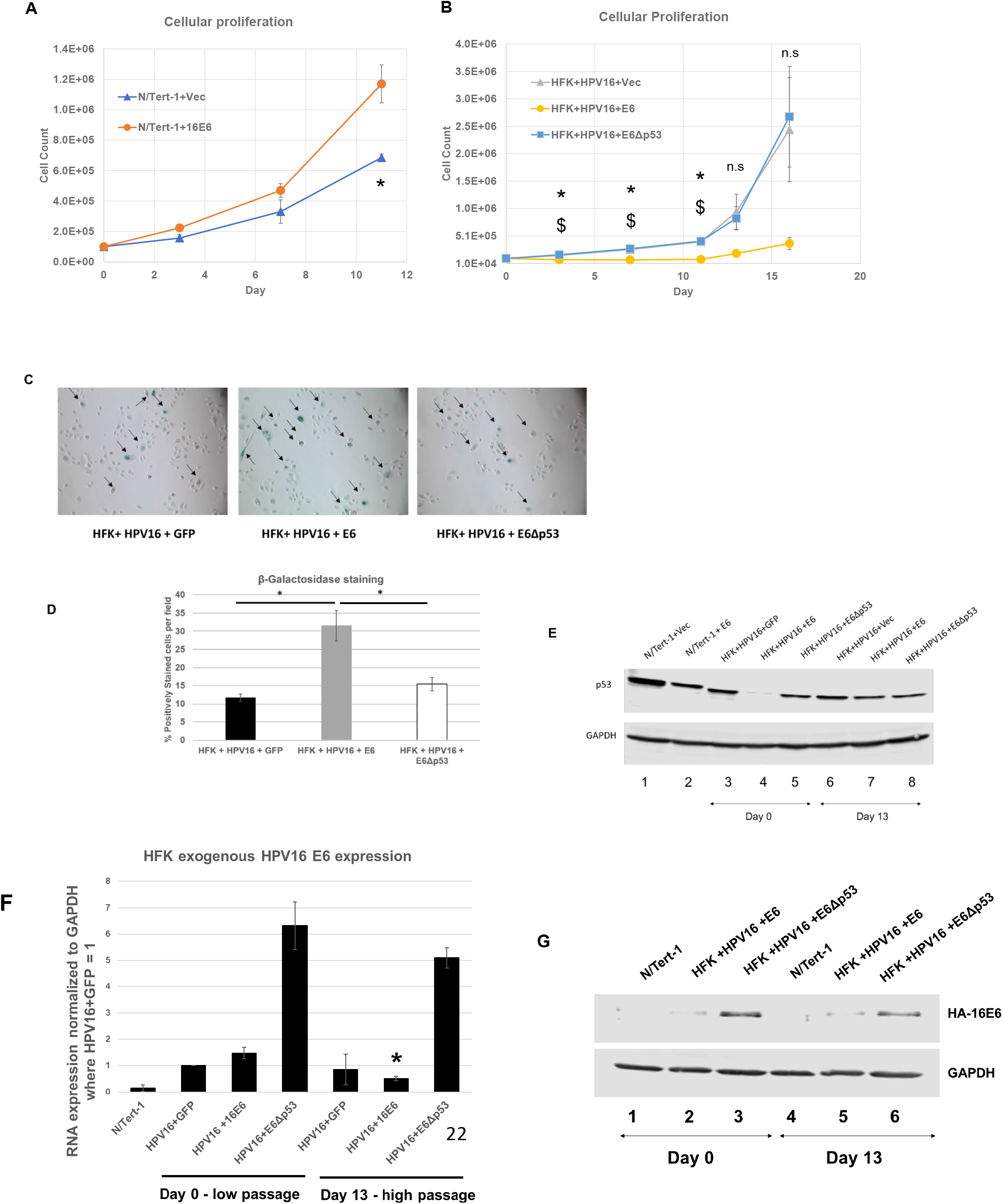
p53 reduction in via introduction of full length HPV16 E6 reduces cellular proliferation in HPV16 immortalized foreskin keratinocytes. (A) 11-day growth curve of N/Tert-1 cells expressing exogenous HPV16 E6 compared to empty vector. (B) 13-day growth curve of human foreskin keratinocytes immortalized by HPV16 and stably expressing exogenous full length E6, mutant E6 that does not bind and degrade p53 (E6Δp53) or GFP control vector. (C) Senescence staining of cells in B at day 11. Arrows indicate positively staining cells. (D) Quantification of senescence staining in C. Western blot analysis of p53 expression following transfection of E6 plasmids (day 0) and after growth rate recovery of HFK+HPV16+E6 (day 13). (E) Western blot analysis of exogenous E6 and E6Δp53 expression at day 0 and day 13 using primary antibody against HA tag. HFK+GFP samples were omitted due to signal oversaturation and bleed over into neighboring wells. GAPDH was used as internal loading control. (F) RT-qPCR analysis of exogenous GFP, E6 and E6Δp53 expression at day 0 and day 13 using primers against FLAG-HA tag. Relative quantity calculated by the ΔΔCT method using GAPDH as an internal control. Bonferroni correction utilized when applicable. (G) Western blotting of the indicated extracts using FLAG antibody (the E6 is double tagged with HA and FLAG).

Overall, these results demonstrate that p53 is expressed in HPV16 immortalized cells, and that this expression may be critical for continuing proliferation of these cells. We next moved on to investigate possible reasons for the requirement of p53 expression in HFK+HPV16 cells.

### Disruption of p53 interaction with the HPV16 E2 protein attenuates cell growth and blocks the viral life cycle

A known and relatively understudied interaction of p53 with HPV16 is the direct physical interaction with E2 (43). We generated an E2 mutant predicted to not interact with p53 (E2^−p53^) and generated stable N/Tert-1 cell lines expressing this mutant, as we have done for wild type E2 (43). There was robust, stable expression of E2 and E2^−p53^ in N/Tert-1 cells (Fig. 3A, lane 3). Immunoprecipitation with a p53 antibody brought down p53, and E2 wild type co-immunoprecipitated with p53 while E2^−p53^ did not (Fig. 3B, compare lanes 2 and 3). To demonstrate that E2^−p53^ was functional we carried out transcriptional studies in N/Tert-1 cells. Because the binding of p53 takes place in the DNA-binding domain (DBD) of E2, we confirmed that the mutant E2 retained DNA-binding function. Both E2 wild type and E2^−p53^ were able to repress transcription from the HPV16 long control region (LCR) efficiently and comparatively (Fig. 3C). We also measured that transcriptional activation function of E2 wild type and E2^−p53^ (Fig. 3D). While E2^−p53^ is able to activate transcription it was compromised when compared with E2 wild type (compare lanes 5-7 with lanes 2-4). We conclude from these experiments that E2^−p53^ is nuclear and able to bind to its DNA target sequences but that its transcriptional activation property (but not repression) is attenuated.

**Figure 3.**
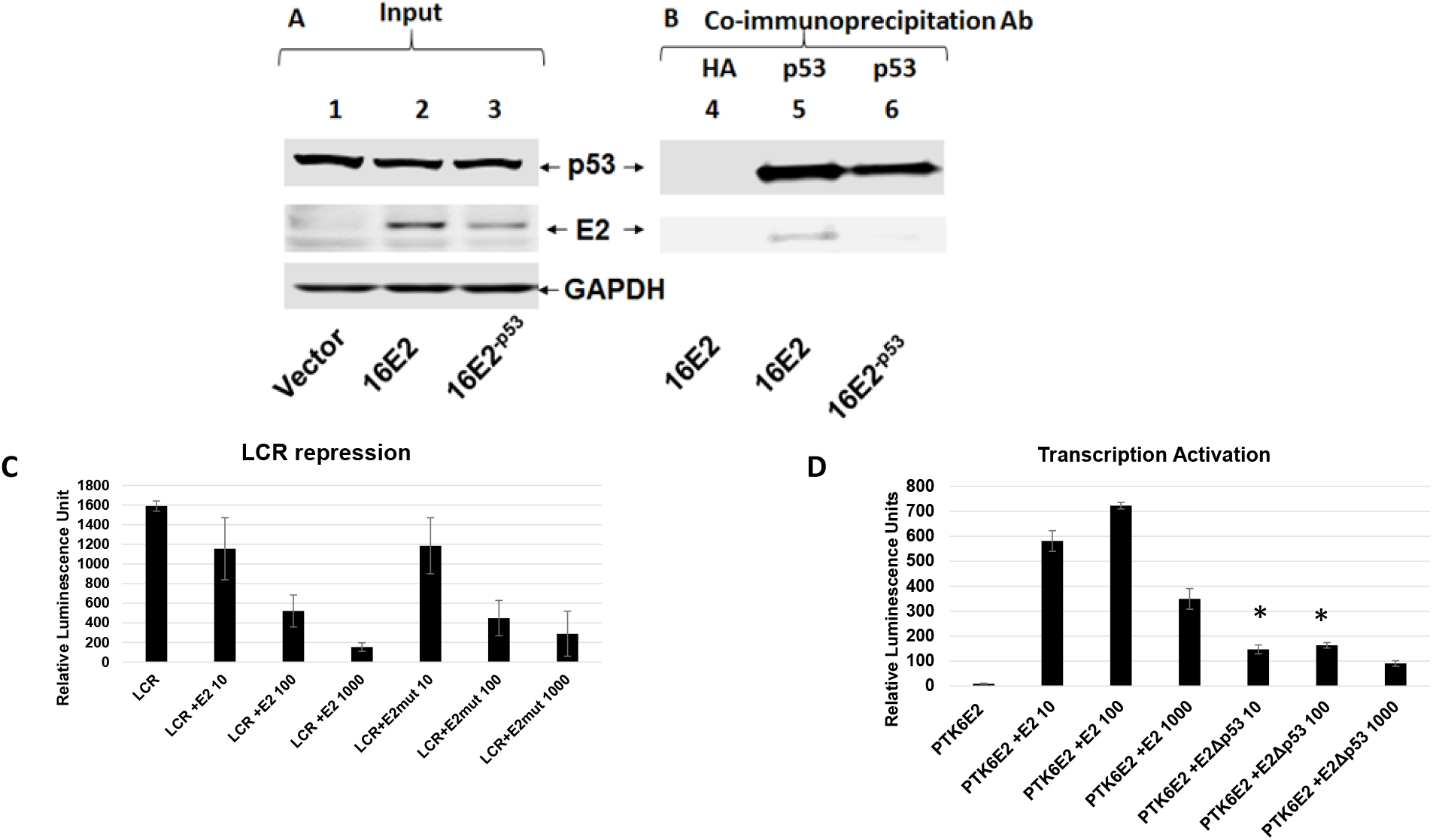
Generation and characterization of p53 binding mutant of HPV16 E2 (E2-p53) in N/Tert-1 cells. (A) Input western blot of stably expressing E2 and E2^−p53^ in N/Tert-1 Cells. For E2^−p53^, residues W341, D344 and D338 were mutated to alanine as previously described (42, 43). (B) Co-immunoprecipitation pull down of E2 using polyclonal antibody against p53. (C) HPV16 long control region repression assay of wild-type E2 and E2^−p53^. N/Tert-1 cells were transiently transfected with 1 μg pHPV16-LCR-Luciferase reporter plasmid along with 10ng, 100ng, or 1000ng of E2 or E2-p53 plasmid. (D) E2 transcriptional activity assay of wild-type E2 and E2-p53. Similar to LCR repression assay, N/Tert-1 cells were transiently transfected with 1 μg pTK6E2-Luciferase reporter plasmid along with increasing amounts of E2 wild-type and E2-p53 plasmids. For (C) and (D), relative luminescence units were calculated by normalizing absolute luminescence readouts to input protein concentration.

Having confirmed that the E2^−p53^ mutant was functional, we introduced the mutations that abrogate the E2-p53 interaction into the entire HPV16 genome (HPV16^−p53^). We introduced the wild type and mutant HPV16 genomes into 2 independent primary human foreskin cell populations that we recently used to investigate the role of the E2-TopBP1 interaction in the viral life cycle (53). Both the wild type and mutant genomes efficiently immortalized both HFK donor cells. We carried out Southern blotting on *Sphl* cut DNA (a single cutter for the HPV16 genome) (Fig. S1A). To further characterize the status of the genomes in these cells we used TV exonuclease assays (this assay is based on the fact that episomal HPV16 genomes are resistant to exonuclease digestion) (54, 55). This assay demonstrated that the viral DNA in the immortalized donor cell lines retained a predominantly episomal status, irrespective of whether the viral genomes were wild type or HPV16^−p53^ (Fig. S1B).

Next, we investigated the expression of markers relevant to HPV infection in HFKs. Fig. 4A demonstrates that p53 levels are similarly reduced in HFK+HPV16 and HFK+HPV16^−p53^ cells when compared with N/Tert-1 cells (compare lanes 2-5 with lane 1) (Fig. 4B). For comparison, cells immortalized with an E6/E7 expression vector had almost no p53 expression (lane 6), likely due to the inability of the E6 to be spliced to E6* variants with this expression vector. To further characterize these cell lines, we investigated whether the DNA damage response is turned on as HPV infections activate both the ATR and ATM pathways. We investigated the phosphorylation status of CHK1 and CHK2 as surrogate markers for activation of these DNA damage response kinases (Fig. 4D). Compared with N/Tert-1 cells there is an overall increase of CHK1 and CHK2 levels in cells immortalized with HFK+HPV16, HFK+HPV16^−p53^ or E6/E7 expression. CHK1 and CHK2 phosphorylation is also elevated in the presence of all of the HPV16 positive cells when compared with N/Tert-1 cells. It is important to note that E6 and E7 immortalization of HFK induced phosphorylation of CHK1 but not CHK2 when compared with the entire genome (Lane 6). This is likely due to the ATM pathway being largely activated by viral replication rather than by the viral oncogenes E6 and E7 which we have previously reported (56). Overall, these results suggest that markers of HPV16 infection are activated in HFK cells immortalized with HPV16 irrespective of the ability of p53 to bind E2.

**Figure 4.**
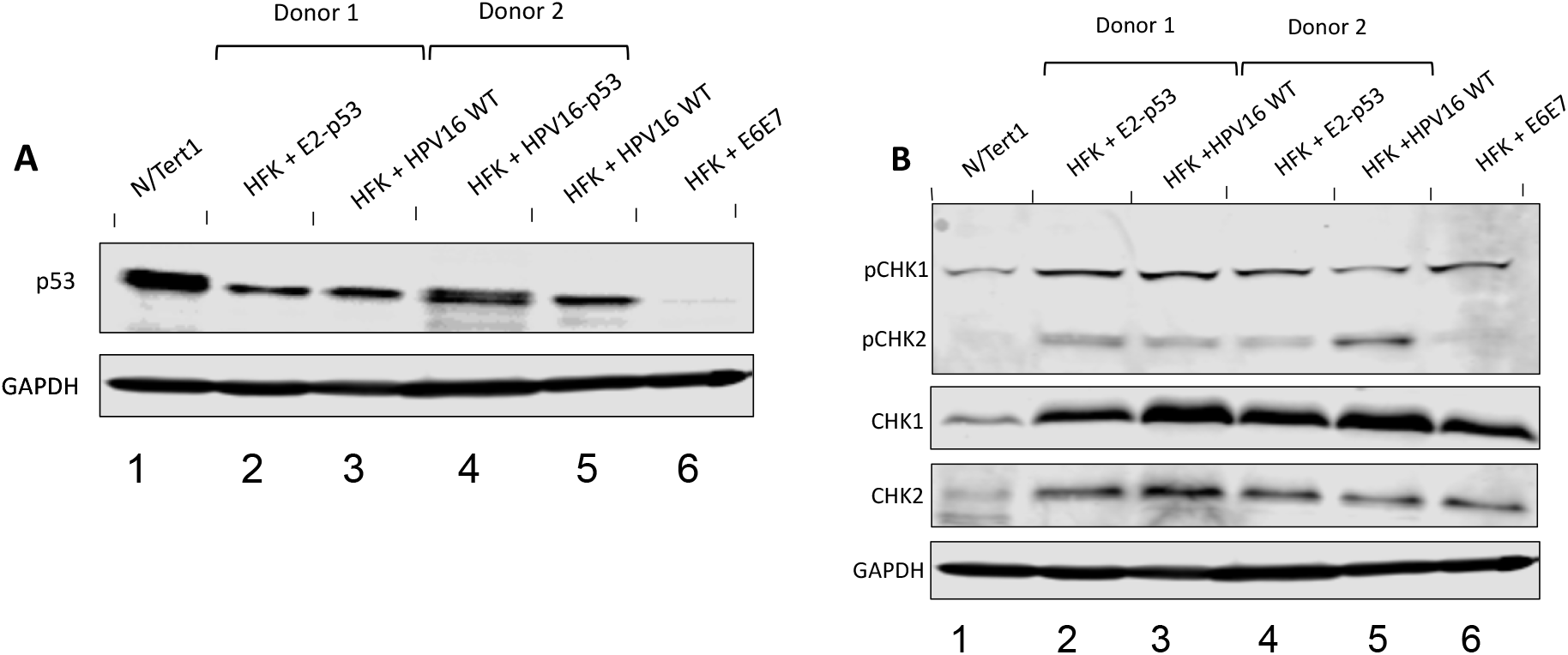
Generation and characterization of HPV16-p53 immortalized human foreskin keratinocytes (HFKs). (A) p53 protein expression in two independent HFK donors immortalized by wild-type HPV16 (Lanes 3 and 5) and HPV-p53 (Lanes 2 and 4). N/Tert-1 and HFK immortalized by E6 and E7 are provided for reference (lanes 1 and 6 respectively). (B) Activation of the ATR and ATM DNA-damage pathways in immortalized HFKs. ATR and ATM activation by HPV16 leads to phosphorylation of Checkpoint kinases 1 and 2 respectively and serve as markers for HPV infection and replication.

Even though the HFK+HPV16^−p53^ cells had markers indicative of HPV16 immortalization, we noticed an aberrant growth phenotype in both foreskin donor cells (Fig. 5A). There was an initial enhanced proliferation of the HFK+HPV16^−p53^ cells when compared with HFK+HPV16. However, around the 3-4 week mark, the HFK+HPV16^−p53^ cells began to slow their growth and eventually stopped proliferating. To determine the mechanism of the attenuation of cell growth we investigated senescence in N/Tert-1, HFK+HPV16 and HFK+HPV16^−p53^ cells by staining for beta-galactosidase (Fig. 5B). There was a significantly increased number of senescent cells with the p53 mutant cells and this was quantitated (Fig. 5C). Senescence can be induced by increased DNA damage, particularly double strand breaks (DSB) (25, 26). Because CHK1 and CHK2 pathway activation was not noticeably different between HFK+HPV16 and HFK+HPV16^−p53^, we decided to look at DSBs more directly using single-cell gel electrophoresis (COMET assay). As expected, the expression of wild type or mutant HPV16 genomes in HFKs led to increased formation of DSBs as indicated by olive tail moment (OTM) when compared to HPV negative N/Tert-1 cells (57) (Fig. 6D). However, the mutant HFKs consistently exhibited larger OTM values compared to HFK+HPV16 (Fig. 5D and 5E). As the expression of full-length E6 from a heterologous vector attenuates the growth of HFK+HPV16 wild type cells (Fig. 2) we rationalized that expression of E6 should not alter the growth of HFK+HPV16^−p53^ cells. Stable expression of exogenous full length E6 or the E6Δp53 mutant had no additional effect on the proliferation of HFK+HPV16^−p53^ cells, illustrating that the drastic differences in proliferation are potentially due to the E2-p53 interaction (Fig. 5F).

**Figure 5.**
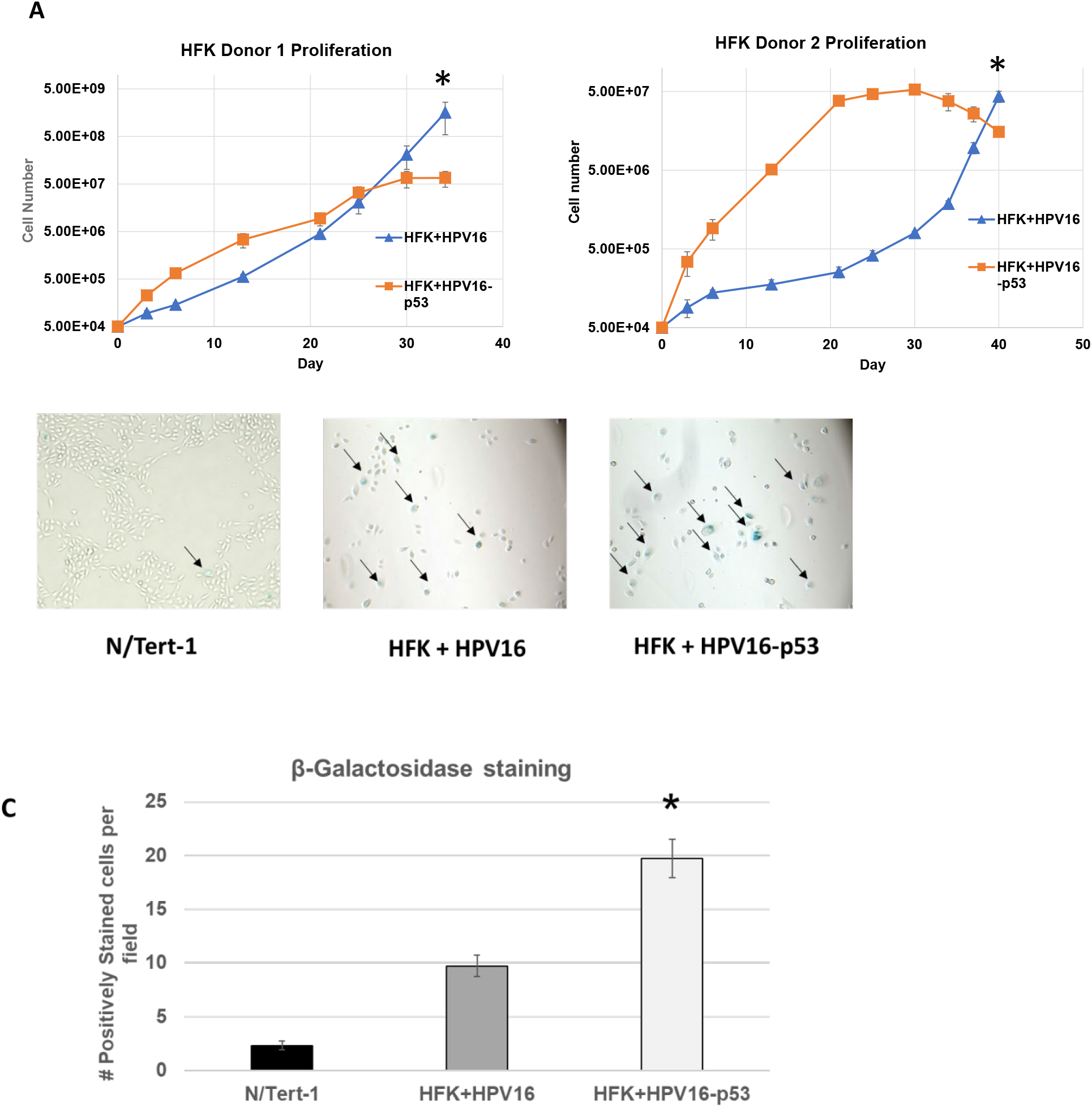
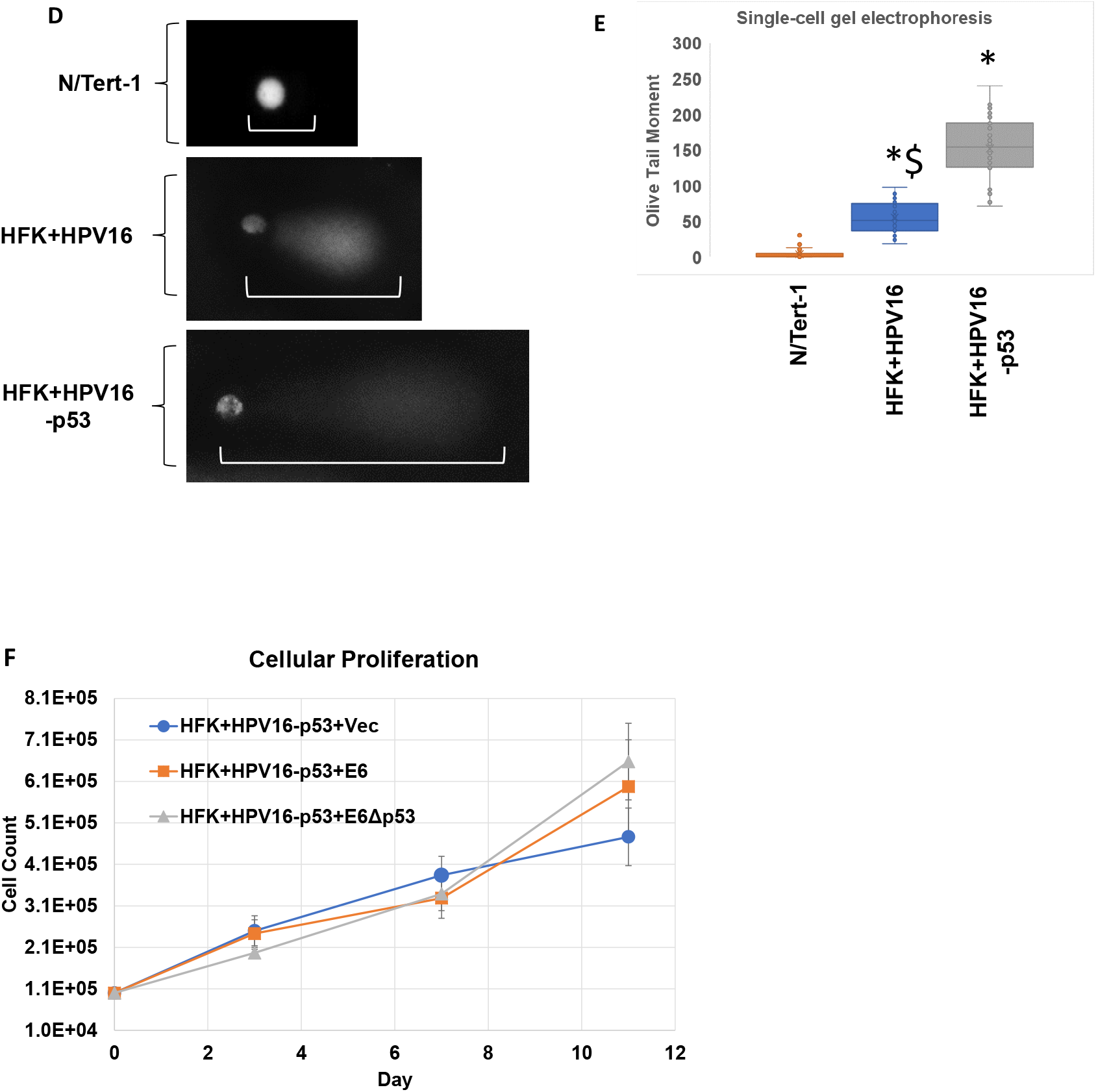
HFKs immortalized by HPV16-p53 exhibit aberrant growth phenotype, elevated levels of senescence and increased DNA damage and fragmentation. (A) Extended growth curve on HFK’s immortalized by wild-type HPV16 and HPV16-p53. Cells were grown over a period of 34-40 days depending on HFK donor cell line. In general, donor 1 proliferated quicker than donor 2 regardless of HPV genome status. (B). β-galactosidase staining as a marker of senescence for proliferating HFK+HPV16 and HFK+HPV16-p53 cells compared to N/Tert-1 cells. Images taken at 10X. Five random fields were imaged per replicate per cell line. Representative image presented with positively stained cells marked by arrows. (C) Quantification of β-galactosidase staining. Average number of positively stained cells per high power field were calculated by a blinded observer +/− SEM. (D) Single-cell gel electrophoresis (COMET) Assay. Cells were grown in 24-well plate for 24 hours then trypsinized, washed, resuspended in 0.5% low molecular weight agarose, and subjected to single cell gel electrophoresis. DNA was stained with DAPI. Five randomly selected fields were imaged at 20x per replicate per cell line. Representative comets are presented with white bars highlighting comet tails. (E) The olive-tail moments (OTMs) of all non-overlapping comets in each high-power field were were quantified using CaspLab COMET assay software. Average OTM +/− SEM. *p<0.05 for HFK+HPV16 vs HFK+HPV16-p53. $p<0.05 for HFK+HPV16 vs N/Tert-1. Bonferoni correction used where applicable. (F). Eleven-day growth curve on HFK+HPV16-p53 stably expressing exogenous E6, E6Δp53 or GFP control.

**Figure 6.**
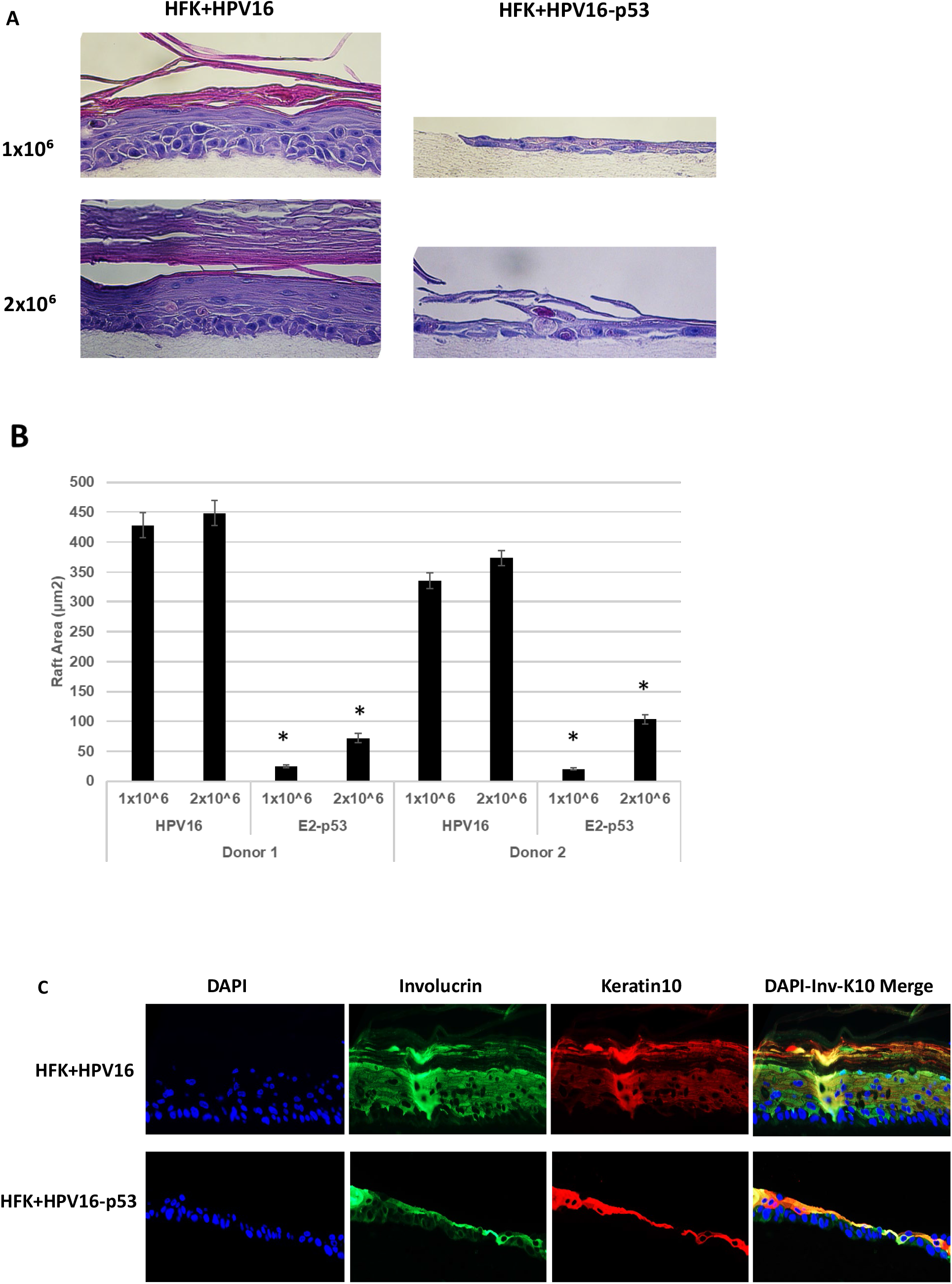
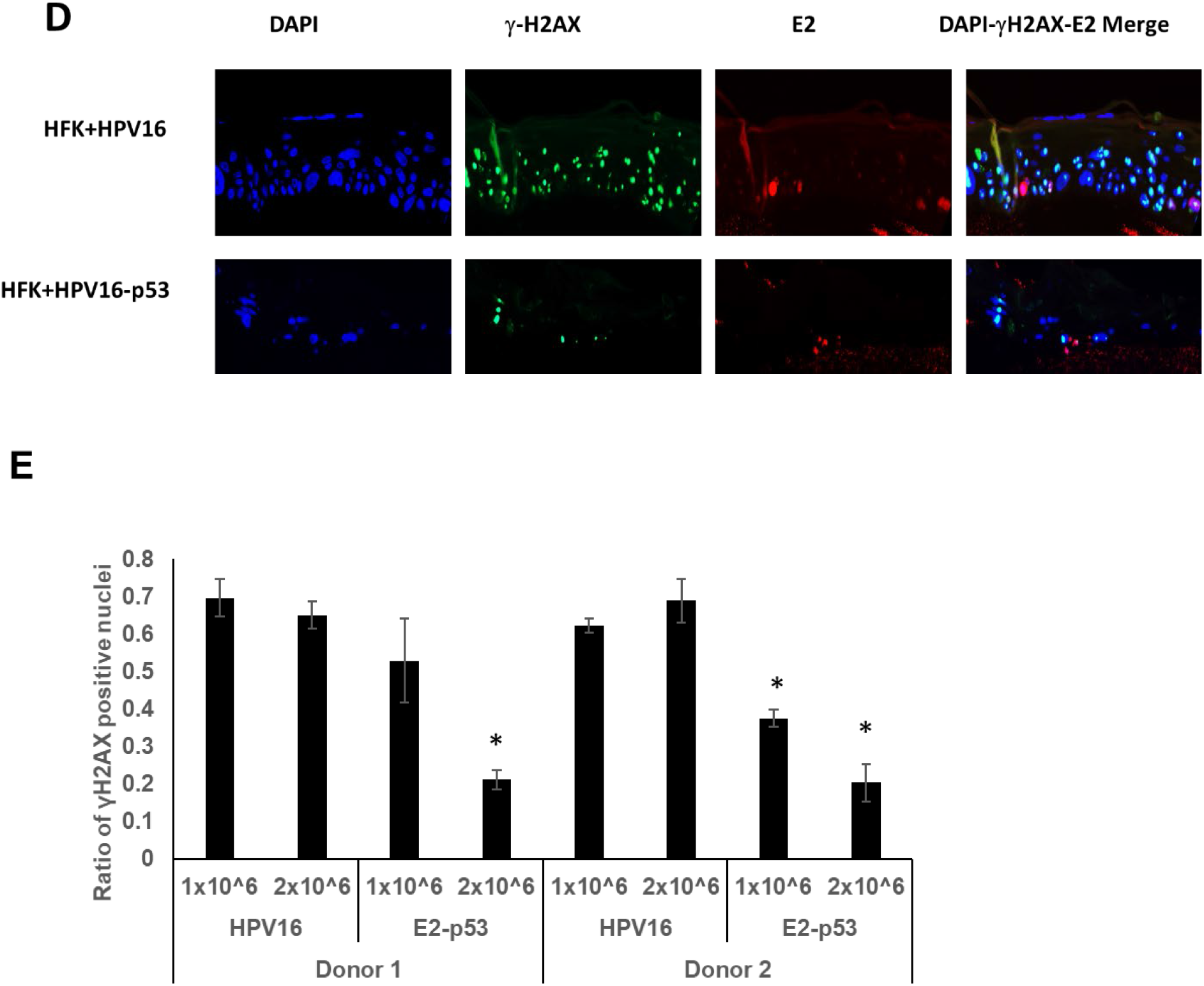
Organotypically rafted HFKs immortalized by HPV16-p53 exhibit aberrant life cycle with dysregulated differentiation, lower markers of viral replication and overall reduced raft proliferation. (A) Organotypic raft cultures and H&E staining of samples from figure 5. HFKs were seeded onto collagen matrices at densities of 1×10^6^ (upper panels) and 2×10^6^ (lower panels). (B). The experiment in A was repeated in a second independent HFK donor and average raft areas were calculated for each donor using a Keyence imaging system. (C) HFK rafts stained using indicated antibodies as markers of keratinocyte differentiation. (D) DNA damage and viral replication marker γ-H2AX was stained for in HPV16 and HPV16-p53 HFK rafts. (E) γ-H2AX staining was repeated in a second HFK donor and quantified using a Keyence imaging system.

### HFK+HPV16^−p53^ cells have an aberrant life cycle in differentiating epithelium

We organotypically rafted HFK+HPV16 and HFK+HPV16^−p53^. Both lines were placed on collagen plugs at early passage when the HFK+HPV16^−p53^ cells retained proliferative capacity. Due to the large difference in growth rates between the wild type and mutant cells, the original plating was performed with both 1×10^6^ and 2×10^6^ cells to promote production of a monolayer on the collagen plugs prior to lifting to the liquid-air interface for differentiation. Fig 6A demonstrates an aberrant differentiation process with the HFK+HPV16^−p53^ cells when compared with HFK+HPV16 cells at both cell densities. It is noticeable that at the lower cell density (1×10^6^) there was a failure to form a monolayer prior to induction of differentiation (as evidenced by gaps between keratinocyte cell clusters on the collagen plug). Using a seeding density of 2×10^6^ eliminated the formation of gaps but did not improve the proliferation. A representative of two independent donors is shown, both donors had identical phenotypes. Fig. 6B quantitates the results from two independent rafts from two independent donors; the mutant genomes have dramatically lower raft area when compared with wild-type genomes. To investigate whether differentiation has occurred in these cells we stained with Involucrin and Keratin 10 (Fig. 6C). The mutant genome cells stained positive for both differentiation markers demonstrating that, even though raft growth is markedly attenuated, differentiation still occurs. We also stained for viral replication using the DNA-damage marker γ-H2AX. Recently we reported that an E2 mutant that failed to interact with TopBP1 results in degradation of E2 during organotypic rafting; this degradation would block viral replication and indeed these cells had no γ-H2AX staining (53). This demonstrates that the γ-H2AX staining indicates the occurrence of viral replication. Fig. 6D demonstrates that there is abundant nuclear γ-H2AX staining throughout HFK+HPV16 cells, indicating replication is occurring. The HFK+HPV16^−p53^ cells also support viral replication although there is a reduction in the number of rafted cells stain positively for γ-H2AX (Fig. 6D).

## Discussion

The HPV E2 protein is essential for viral genome replication, segregation of viral episomes into daughter cells following cell division, and can transcriptionally regulate both virus and host genomes (12, 19, 20). E2 interacts with a variety of host factors to promote progression of the viral life cycle, many of which are essential such as interactions with TopBP1 and BRD4 (12, 19, 23, 53, 58). In this report, we propose that E2 binding to p53 is also an essential interaction as abrogation of the interaction leads to catastrophic failure of the viral life cycle.

In the classical high-risk HPV model, upon initial infection, the viral oncogenes E6 and E7 inhibit and degrade tumor suppressor proteins p53 and pRb respectively, promoting hyperproliferation, unregulated DNA replication, mutation accumulation and potentially eventual carcinogenesis. Therefore, immortalization of cell lines can be achieved with overexpression vectors of E6 and E7 (Fig. 1A, Lanes 4 and 8). Previous studies suggest that E6 splice variants and their action on E6-E6AP-p53 complex disruption is cell-cycle dependent (38). In HPV18 cell lines, E6*I shows marked upregulation during and p53 restoration during G2/M (38). We have previously illustrated that E2 is stabilized during mitosis which is important for its association with TopBP1 and its role as a segregation factor (53). It is entirely possible that the cell-cycle mediated p53 restoration corresponds with E2 stabilization allowing these proteins to interact, and may also play an important role in genome segregation.

E2 can regulate host transcription in a multitude of ways. We recently reported that E2 can epigenetically repress the TWIST1 at the histone level, inhibiting EMT and promoting a less aggressive cellular phenotype (20). E2 can also promote the recruitment of DNA methyltransferase 1 to interferon response genes, resulting in DNA base methylation and global innate immunity downregulation (21). It is currently unclear how E2 recruits epigenetic modifiers to these genes and p53 may play an important role. DNA methyltransferases (DMNTs) are often part of large multimeric complexes and use transcription regulatory proteins to help target specific genes undergoing epigenetic silencing (59, 60). p53 is known to also interact with DMNT1 resulting in the methylation of antiapoptotic genes (61). It is possible that the interaction between E2 and p53 is important for the rerouting of DNMTs to different genes whose regulation is important for a healthy viral life cycle. It is also noticeable that E2-p53 has an attenuated ability to activate transcription (Fig. 3), indicating that regulation of host gene transcription by E2 may require co-operation with p53 in some cases.

The results from Fig. 5D suggest that additional double strand breaks play a role in the enhanced damage and proliferation rate of HFK+HPV16-p53 mutant cells compared to wild type immortalized HFKs. HPV uses homologous recombination (HR) factors to assist in viral replication (62–64). Conversely, p53 binds to replication protein A (rpA) resulting in repression of HR, reducing DSB repair and promoting apoptosis during catastrophic genome instability (27, 29). It is possible that E2 helps regulate this activity of p53 and inability to do so results in accumulation of DSBs as seen in Fig. 5D.

In conclusion, this report indicates that p53 expression is retained in HPV16 positive cell lines and tumor samples under a variety of conditions. Depletion of this residual p53 by full length E6 results in significant reduction in proliferation and enhanced senescence in cells immortalized with HPV16 which we attribute to loss of E2 interaction with p53. Human foreskin cells immortalized by HPV16 where E2 can no longer bind to p53 exhibit aberrant phenotypes including dysregulated proliferation, enhanced levels of DSBs and overall failure of the viral life cycle during organotypic raft culturing. Due to the importance of p53 in the context of HPV related cancers as well as the profound phenotypes demonstrated in this report, further investigation on the interaction between E2 and p53 is warranted.

## Methods

### Cell culture

N/Tert-1 cells and head-and-neck cancer lines UMSCC47 and UMSCC104 were cultured as previously described (20, 21, 65–67). Immortalization and culturing of human foreskin keratinocytes with HPV16 are described below. Cells were incubated at 37°C and 5% CO2 with media changed every 3 days. For cisplatin treatment, cells were incubated with indicated concentrations of drug dissolved in DMF or DMF vehicle control for 24-hours

### Immortalization of human foreskin keratinocytes (HFK)

HPV16 mutant genome (HPV16-p53, which contained an E2 unable to bind p53) was generated and sequenced by Genscript (42, 44, 68). Residues Aspartic acid 388, Tryptophan 341 and Aspartic acid 344 were all mutated to alanine resulting in inability for E2 to interact with p53. The HPV16 genome was removed from the parental plasmid using *Sphl*, and the viral genomes isolated and then re-circularized using T4 ligase (NEB) and transfected into early passage HFK from three donor backgrounds (Lifeline technology), alongside a G418 resistance plasmid, pcDNA. Cells underwent selection in 200 μg/mL G418 (Sigma-Aldrich) for 14 days and were cultured on a layer of J2 3T3 fibroblast feeders (NIH), which had been pre-treated with 8 μg/ml mitomycin C (Roche). Throughout the immortalization process, HFK were cultured in Dermalife-K complete media (Lifeline Technology).

### Western blotting

Protein from cell pellets was extracted with 2x pellet volume protein lysis buffer (0.5% Nonidet P-40, 50mM Tris [pH 7.8], and 150mM NaCl) supplemented with protease inhibitor (Roche Molecular Biochemicals) and phosphatase inhibitor cocktail (Sigma). Protein extraction from patient derived xenografts was performed as previously described.(45, 46) The cells were lysed on ice for 30 min followed by centrifugation at 18,000 rcf (relative centrifugal force) for 20 min at 4°C. Protein concentration was estimated colorimetrically using a Bio-Rad protein assay and 25 μg of protein with equal volume of 2X Laemmli sample buffer (Bio-Rad) was denatured at 70°C for 10 min. The samples were run on a Novex™ WedgeWell™ 4 to 12% Tris-glycine gel (Invitrogen) and transferred onto a nitrocellulose membrane (Bio-Rad) using the wet-blot method, at 30 V overnight. The membrane was blocked with *Li-Cor* Odyssey® blocking buffer (PBS) diluted 1:1 v/v with PBS for 1 hour at room temperature and then incubated with specified primary antibody in *Li-Cor* Odyssey® blocking buffer (PBS) diluted 1:1 with PBS. Afterwards, the membrane was washed with PBS supplemented with 0.1% Tween20 and further probed with the Odyssey secondary antibodies (IRDye® 680RD Goat anti-Rabbit IgG (H + L), 0.1 mg or IRDye® 800CW Goat anti-Mouse IgG (H + L), 0.1 mg) in *Li-Cor* Odyssey® blocking buffer (PBS) diluted 1:1 with PBS at 1:10,000 for 1 hour at room temperature where applicable. After washing with PBS-tween, the membrane was imaged using the Odyssey^®^ CLx Imaging System and ImageJ was used for quantification. Primary antibodies used for western blotting studies are as follows: p53, 1:1000 (Santa Cruz; cat. no. sc-47698) HPV16 E2, 1:1000 (TVG261) (Abcam; cat. no. ab17185) phospho-CHK1, 1:1000 (Ser345) (Cell Signaling; cat. No. 2341S), phosphor CHK2 (Thr68), 1:1000 (Cell Signaling; cat. No. 2661S), CHK1, 1:1000(Cell Signaling; cat. No. 2360), CHK2, 1:1000 (Abcam; cat. No. ab47443), GAPDH, 1:250 (Santa Cruz; cat. no. sc-47724).

### Plasmids

The following plasmids were used the completion of these studies: pMSCV-N-FLAG-HA-GFP, pMSCV-N-FLAG-HA-HPV16E6, pMSCV-IP-N-FLAG-HA-16E6 8S9A10T (“E6Δp53”). Wild-type 16E2 or 16E2-p53 (Mutated residues W341A, D344A, D338A) were cloned into pCDNA vector for confirmation of p53 interaction in N/Tert-1 cells. pCDNA was used for empty vector control.

### Real-time qPCR

RNA was isolated using the SV Total RNA isolation system (Promega) according to manufacturer’s instructions. 2 μg of RNA was reverse transcribed into cDNA using the high-capacity reverse transcription kit (Applied Biosystems). The PowerUp SYBR green master mix (Applied Biosystems) was used along with cDNA and gene specific primers and real-time PCR was performed using a 7500 Fast real-time PCR system as previously described. (20, 21, 65) Expression was quantified as relative quantity over GAPDH using the 2^−ΔΔC_T_^ method. Primers used are as follows. FLAG-HA Tag fwd 5’- GACTACAAGGATGACGATG- 3’, FLAG-HA Tag rev 5’- GCGTAATCTGGAACATCG −3’.

### p53 Immunoprecipitation

Primary polyclonal antibody against p53 (Invitrogen; PA5-27822) or a HA-tag antibody (used as a negative control) was incubated in 200 μg of cell lysate (prepared as described above), made up to a total volume of 500 μl with lysis buffer (0.5% Nonidet P-40, 50mM Tris [pH 7.8], and 150mM NaCl), supplemented with protease inhibitor (Roche Molecular Biochemicals) and phosphatase inhibitor cocktail (Sigma) and rotated at 4°C overnight. The following day, 50 μl of pre-washed protein A-sepharose beads per sample was added to the lysate/antibody solution and rotated for 4 hours at 4°C. The samples were gently washed with 500 μl lysis buffer by centrifugation at 1,000 rcf for 2-3 min. This wash was repeated 4 times. The bead pellet was resuspended in 4X Laemmli sample buffer (Bio-Rad), heat denatured and centrifuged at 1,000 rcf for 2-3 min. Proteins were separated using an SDS-PAGE system and transferred onto a nitrocellulose membrane before probing for the presence of E2 or p53, as per western blotting protocol.

### Transcription and LCR Repression assays

A pTK6E2-Luciferace reporter plasmid was utilized to analyze transcriptional activation of Wild-type HPV16 E2 and E2Δp53 proteins as previously described (23). 5×10^5^ N/Tert-1 cells were seeded onto 100mm^2^ plate and transfected 24 hours later with 0 ng, 10 ng, 100 ng or 1000 ng of E2 WT or E2^−p53^ plasmid DNA along with 1000ng of pTK6E2-Luciferase reporter plasmid using Lipofectamine 2000 according to the manufacturer’s instructions (ThermoFisher Scientific). Cells were harvested the next day using Promega luciferase assay system. For LCR repression activity, a pHPV16-LCR-Luciferase reporter was used in place of the pTK6E2-Luciferase plasmid.(65)

### Southern blotting

Total cellular DNA was extracted by proteinase K-sodium dodecyl sulfate digestion followed by a phenol-chloroform extraction method. 5 μg of total cellular DNA was digested with either SphI (to linearize the HPV16 genome) or HindIII (which fails to cut HPV16 genome). All digestions included DpnI to ensure that all input DNA was digested. All restriction enzymes were purchased from NEB and utilized as per manufacturer’s instructions. Digested DNA was separated by electrophoresis of a 0.8% agarose gel, transferred to a nitrocellulose membrane, and probed with radiolabeled (32-P) HPV16 genome as previously described.(67) This was then visualized by exposure to film for 1 to 24 hours. Images were captured from an overnight-exposed phosphor screen by GE Typhoon 9410 and quantified using ImageJ.

### Exonuclease V assay

PCR based analysis of viral genome status was performed using methods described by Myers et al (2019). Briefly, 20 ng genomic DNA was either treated with exonuclease V (RecBCD, NEB), in a total volume of 30 μl, or left untreated for 1 hour at 37°C followed by heat inactivation at 95°C for 10 minutes. 2 ng of digested/undigested DNA was then quantified by real time PCR using a 7500 FAST Applied Biosystems thermocycler with SYBR Green PCR Master Mix (Applied Biosystems) and 100 nM of primer in a 20 μl reaction. Nuclease free water was used in place of the template for a negative control. The following cycling conditions were used: 50°C for 2 minutes, 95°C for 10 minutes, 40 cycles at 95°C for 15 seconds, and a dissociation stage of 95°C for 15 seconds, 60°C for 1 minute, 95°C for 15 seconds, and 60°C for 15 seconds. Separate PCR reactions were performed to amplify HPV16 E6 F: 5’- TTGCTTTTCGGGATTTATGC-3’ R: 5’- CAGGACACAGTGGCTTTTGA-3’, HPV16 E2 F: 5’- TGGAAGTGCAGTTTGATGGA-3’ R: 5’- CCGCATGAACTTCCCATACT-3’, human mitochondrial DNA F: 5’- CAGGAGTAGGAGAGAGGGAGGTAAG-3’ R: 5’-TACCCATCATAATCGGAGGCTTTGG −3’, and human GAPDH DNA F: 5’- GGAGCGAGATCCCTCCAAAAT-3’ R: 5’- GGCTGTTGTCATACTTCTCATGG-3’

### Senescence Staining

7.5×10^4^ cells were seeded in 6-well plates. The following day, cells were stained for senescence using the Cell Signal Senescence β-Galactosidase Staining kit according to manufacturer’s instructions (9860). Randomly selected images were taken using the Keyence imaging system at 10X. Positively stained cells were counted by a blinded observer and average number of positively stained cells per field were calculated.

### Single-cell gel electrophoresis (COMET) Assay

1×10^4^ cells were plated in 24-well plate with 1mL media one day prior to harvest. The next day, cells were trypsinized and resuspended in a mixture 0.5% w/v Low molecular weight agarose (Lonza, cat. No. #50101) and PBS at a ratio of 10:1. Suspension was immediately pipetted onto Trivegen COMET Slides™ (4250-004-03) and allowed to dry for 30 min at 4°C. Slides underwent lysis for 90 min at 4°C in the dark (Lysis buffer: 10mM Tris, 100mM EDTA, 2.5M NaCl, 1% TritonX100, 10%DMSO titrated to pH 10.0). Afterwards slides were placed in Alkaline buffer for 25 min at at 4°C in the dark (Alkaline buffer: 1mM EDTA, 200mM NaOH, pH >13.0). Slides were transferred to an agarose gel electrophoresis box filled with additional alkaline buffer. Electrophoresis was performed at 25V for 20min at room temperature in the dark. Slides were then washed 2x in dd (double distilled) H2O for 5 min at RT and then placed in neutralization buffer for 20min at RT in dark (Neutralization buffer: 400mM Tris-HCl titrated to pH 7.5). Neutralized slides were then left to dry at 37°C in the dark. Dried slides were stained with DAPI (1:10,000 in dd H2O) for 15 min at RT then washed 2x with dd H20 for 5 min. Stained and rinsed slides were left to dry overnight. Slides were imaged using the Keyence imaging system at 20x with >5 images taken per replicate. Quantization of olive tail moments (OTM) was achieved using the CASPLab COMET Assay imaging software by Końca K et al, 2003 (69).

### Organotypic raft culture

Keratinocytes were differentiated via organotypic raft culture as described previously.(21, 67, 70) Briefly, cells were seeded onto type 1 collagen matrices containing J2 3T3 fibroblast feeder cells. Cells were cultured to confluency atop the collagen plugs, lifted onto wire grids and cultured in cell culture dishes at the air-liquid interface. Media was replaced on alternating days. Following 14 days of culture, rafted samples were fixed with formaldehyde (4% v/v) and embedded in paraffin. Multiple 4μm sections were cut from each sample. Sections were stained with hematoxylin and eosin (H&E) and others prepared for immunofluorescent staining via HIER. Fixing and embedding services in support of the research project were generated by the VCU Massey Cancer Center Cancer Mouse Model Shared Resource, supported, in part, with funding from NIH-NCI Cancer Center Support Grant P30 CA016059. Fixed sections were antigen retrieved in citrate buffer, and probed with the following antibodies for immunofluorescent anaylsis: phospho-yH2AX 1/500 (Cell Signaling Technology; 9718), Involucrin 1/1000 (abcam; ab27495), Keratin 10 1/1000 (SigmaAldrich; SAB4501656), and HPV16 E2. (monoclonal B9)(71) Cellular DNA was stained with 4’,6-diamidino-2-phenylindole (DAPI, Santa Cruz sc-3598). Microscopy was performed using the Keyence imaging system,

## Acknowledgements

This work was supported by VCU Philips Institute for Oral Health Research and the National Cancer Institute designated Massey Cancer Center grant P30 CA016059 (IMM). DB is supported by R01DE027185, a grant from the National Institute of Dental and Craniofacial Research.

